# Morphological evaluation of rat cerebellum following administration of nitrogen monoxide precursor and inhibitor

**DOI:** 10.1101/2022.07.22.501134

**Authors:** Hamidreza Chegini, Azin Chegini, Amir Gholamzad, Hadis Sadeghi, Seyed Mohammad Hossein Noori Mougahi

## Abstract

In this study, the quantitative and qualitative effects of nitrogen monoxide on cerebellar histopathology, by increasing or decreasing in vivo production of this substance, were investigated. In this study, forty Wistar female rats (RAT) with a weight of about 200 to 250 grams and an average age of eight weeks were used. Rats were divided into five groups of eight, including control groups, normal saline, L-NAME, L-Arginine, L-NAME + L-Arginine. On the third, fourth and fifth days, the injection was performed intraperitoneally and on the eighteenth day, after anesthesia with ether and then craniotomy, the brain and cerebellum of the animals were removed and After quantitative measurements including weight and volume, organs were fixed in 10% formalin and after tissue preparation steps, to prepare the slide, sections with a thickness of 5 to 6 microns were prepared from the samples and by general method such as Hematoxylin-eosin and special techniques like Mason trichrome and toluidine blue were stained and evaluated. The results of this study show that in the case of cerebellum, there is no significant difference in the quantitative weight parameter between the control groups, normal saline, L-Arg, L-NAME and L-NAME + L-Arg groups. Regarding the volume parameter, a significant increase (P <0.01) was observed in L-Arg group compared to L-NAME, L-Arg + L-NAME and normal saline groups. In microscopic qualitative parameters, the most changes in the L-Arg group were seen as follows: The granular and molecular layers of the cerebellum became slightly thickened, some of the nuclei in the granular and molecular layers became severely hyperchromatized, and Purkinje cell accumulation was seen with lymphocytic invasion. In the other groups, there were no significant changes. It is inferred from this study that L-Arginine can cause histopathological changes in cerebellar tissue by increasing NO levels in cerebellum. However, in this study, unlike other similar studies, L-NAME injection did not cause significant histopathological change.

## Introduction

Nitric oxide (NO) is a simple diatomic molecule which has a wide biological effects and is produced by different types of cells[1]. Nitric oxide synthase (NOS) (a NO-producing enzyme from L-Arginine) uses molecular oxygen, NADPH and other cofactors. NOS has three isoforms: nNOS (neuronal), iNSO (inducible) and eNOS (endothelial) The latter type is present in macrophage / monocyte cells, fibroblasts, smooth muscle cells and small vessel endothelial cells, heart cells, liver cells and megakaryocytes.[1,2]

The function of the nNOS-containing nerve cells is not determined well however nNOS is likely to cause neurogenic vasodilation. In the central nervous system, nNOS could also be an important local regulator of cerebral circulation, In addition, NO is responsible for many physiological functions of the brain, such as acute mediation of neural behaviors. [2]

Derived NO from endothelial cell’s eNOS is also an important factor for cerebral blood flow. In cardiovascular system[1,3], NO is responsible for controlling blood pressure by relaxing smooth muscle and vasodilation.[4]

In neurons and glial cells, nNOS is involved in regulating cerebrovascular blood flow, memory and learning (through its long term effects on the CNS).

cerebellar oligodendrocytes are a huge target for Nnos physiological signals along with increasing their growth and maturation. In addition NO can also mediate the myelination of axons during evolution. [5] On the other hand, permanent inhibition of nitric oxide production, induces apoptosis of cerebellar granule cells which confirms the role of NO in the survival of granular neurons. [6]

NO is also a potential mediator of non-adrenergic-non-cholinergic (NANC) neurons and therefore could regulate the cardiac contractility and heart rate, gastrointestinal motility, bronchial tone, and erection.[1,3]

nitric oxide is a lipophilic free radical so it can easily cross cell membranes [7]. Therefore, it can be assumed that nitric oxide is present throughout the cerebral tissue, but nitric oxide is an unstable molecule in the body and is immediately converted to stable compounds such as nitrite and nitrate [8]. NO-induced cytotoxicity, appears to be important in the central nervous system. As mentioned, NO is a free radical that can cause tissue damage if overproduced and is involved in many neurodegenerative diseases [4]. Following neurogenic lesions, large amounts of L-glutamate are released, causing neurotoxicity by activating NMDA receptors. Activation of these receptors increases the activity of nNOS. Released nitric oxide, has a toxic effect on adjacent cells, whereas nNOS-containing cells themselves are resistant to the toxic effects of the receptor stimulation. NO, on the other hand, may have protective effects on central nervous system, which is exerted by oxidation-reduction (redox) reaction. NO + reacts with the NMDA receptor and preventing excessive Ca ++ entry into the cellular cytosol. However, NO itself increases the entry of excessive Ca ++ into the cellular cytosol which is leading to toxic effects on the cell [1]. Excessive amounts of NO, on the other hand, are toxic to neurons due to greater response of glutamate receptors (NMDAs) which allow entering large amounts of Ca ++, k +, and Na + to the cell. This condition has significant toxic effects on nerve cells because of excessive intracellular calcium, free radicals and proteases [9]. Therefore, both protective and destructive effects for NO on the central nervous system have been reported. By examining the effects of NO, we will be able to identify the changes in the histopathology of the cerebral cortex, hippocampus and cerebellum and the desired clinical results can be achieved by prescribing inhibitors or inducers of its production.

## Materials and Methods

200 mg of L-arginine powder and 20 mg of L-NAME powder (per kg) were dissolved in normal saline and injected into mice on days 3, 4 and 5. Disposable syringes were used for each injection and Drugs were refrigerated at injection intervals. The intraperitoneal method was used to prescribe drugs, as it’s been used in most similar studies. For dissection of mice On the 18^th^ day, each mouse was anesthetized with pentobarbital and then the brain was removed. After organs removal, mice were killed by chloroform solution. For measuring the volume of indeterminate spatial shape, we use a graduated cylinder and a liquid with a specific volumetric mass such as water, also a scale with an accuracy of one hundredth of a gram was used to the weight of tissues.

### Preparing tissue for microscopic examination

#### Fixation

To prevent post-mortem changes, precipitating cell proteins with solutions such as 10% formalin suggested.

#### Passaging the tissues

during four stages: Dehydrating, Clarification, Infiltration and Embedding, the required strength and hardness are given to the tissue for cutting with a microtome device.

For dehydration, the samples were passed through 70, 80, 96 and absolute alcohols. Xylene is used as a clearing agent to remove alcohol.

For dehydration, the samples are passed through alcohols with concentrations of 70, 80, 96 and absolute alcohol. Xylene is used as a clearing agent to remove alcohol. For impregnation and molding, the tissues are placed in the middle of molds and they are inserted into molten paraffin. After the paraffin has frozen, the samples are ready for cutting.

#### Cutting and fixing the incisions on the slide

Incisions with a thickness of 1 to 2 microns were prepared from the molds and fixed on a slide by a microtome machine, then paraffin is removed.

#### Staining

Hematoxylin-eosin staining (H & E)

This type of staining is the most common one. Hematoxylin is mainly used to show nuclei and cytoplasmic basophilic inclusions. Eosin as the opposite color, ?? As a result, by this method, the cell nucleus becomes blue and the cytoplasm, RBCs and connective tissue red.

#### Toluidine blue staining

This is a blue-based dye, which is used for nucleus. This method used for metachromatic dye of special structure (such as mast cells containing heparin granules or a cartilage matrix rich in chondroitin sulfate). It’s also used for RNAase, RNA and mucopolysaccharide staining.

#### Masson trichrome staining

This material makes the connective tissue visible. In this staining, in comparison with Wengison dye, tungsten phosphomolybdic acids are used as an antifouling agent (stabilizer), along with a hematoxylin dye (Wiggert Iron). As a result of this staining, the cell nucleus is black, the cytoplasm and muscle and creatine are red, and the collagen and mucus are blue or green [15].

As a result, the nucleus can be seen in black, the cytoplasm and muscle, and creatine in red, and collagen and mucus in blue or green [15].

#### Mounting

The stained sections are glued to the slide using the Canada balsam and a microscopic slide is placed on it.

#### Data analysis

SPSS statistical software is used to analyze the data. we also used One way ANOVA to determine the relationship between groups we used One way ANOVA. A significance level of P <0.05 is considered for all comparisons.

## Results

### Quantitative macroscopic results of cerebellar weight

The results of comparing the weight and volume of the cerebellum for each group are given in the tables and graphs.

### Cerebellum weight

To determine the relationship between the groups, one way ANOVA was used which did not show a statistically significant difference (P <0.01) between the groups.

**Table 4a:**
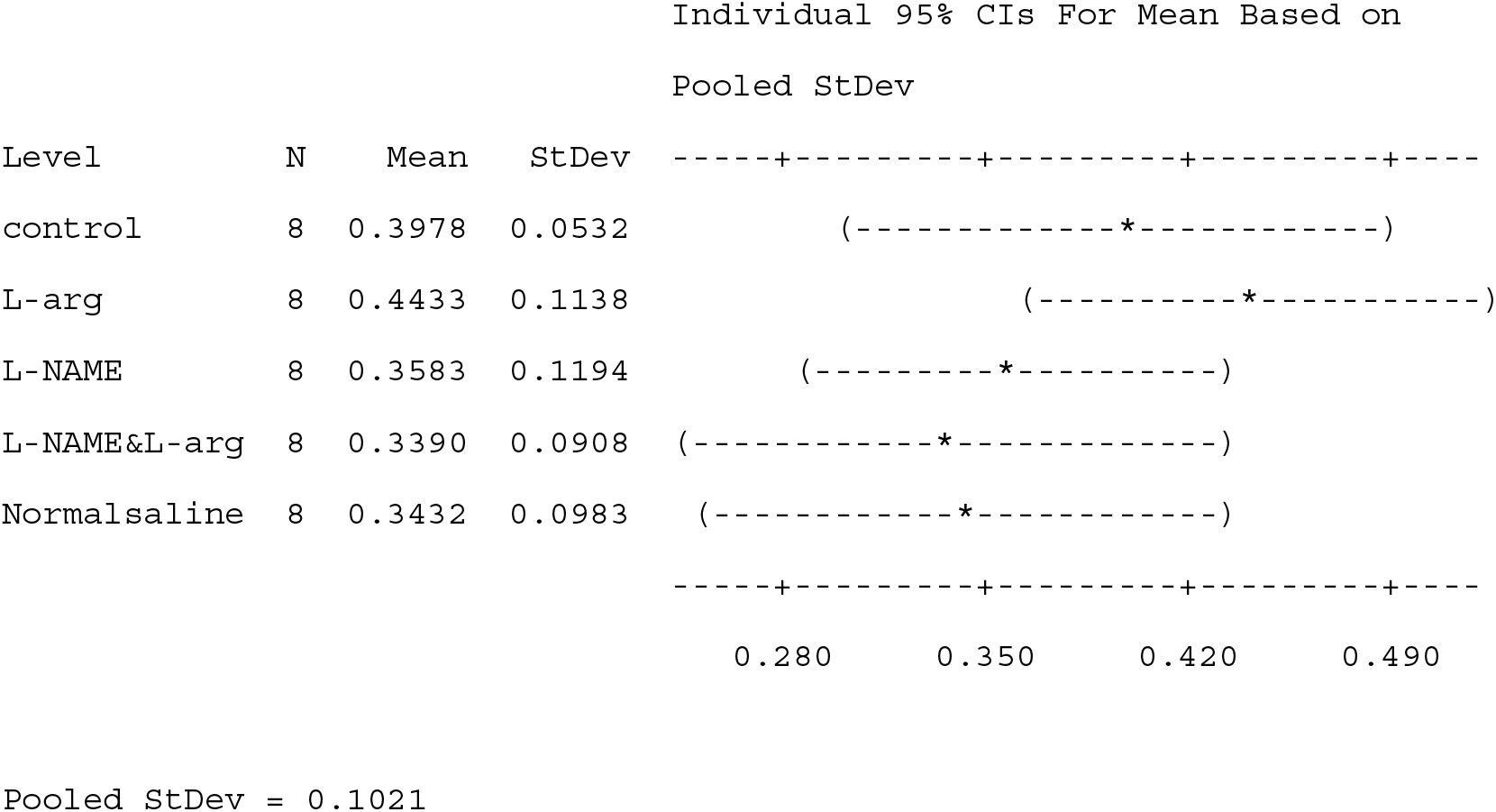
Quantitative values include mean and standard deviation of cerebellar weight in different groups. To the right is the 95% confidence interval of the mean.

**Table 4b:**
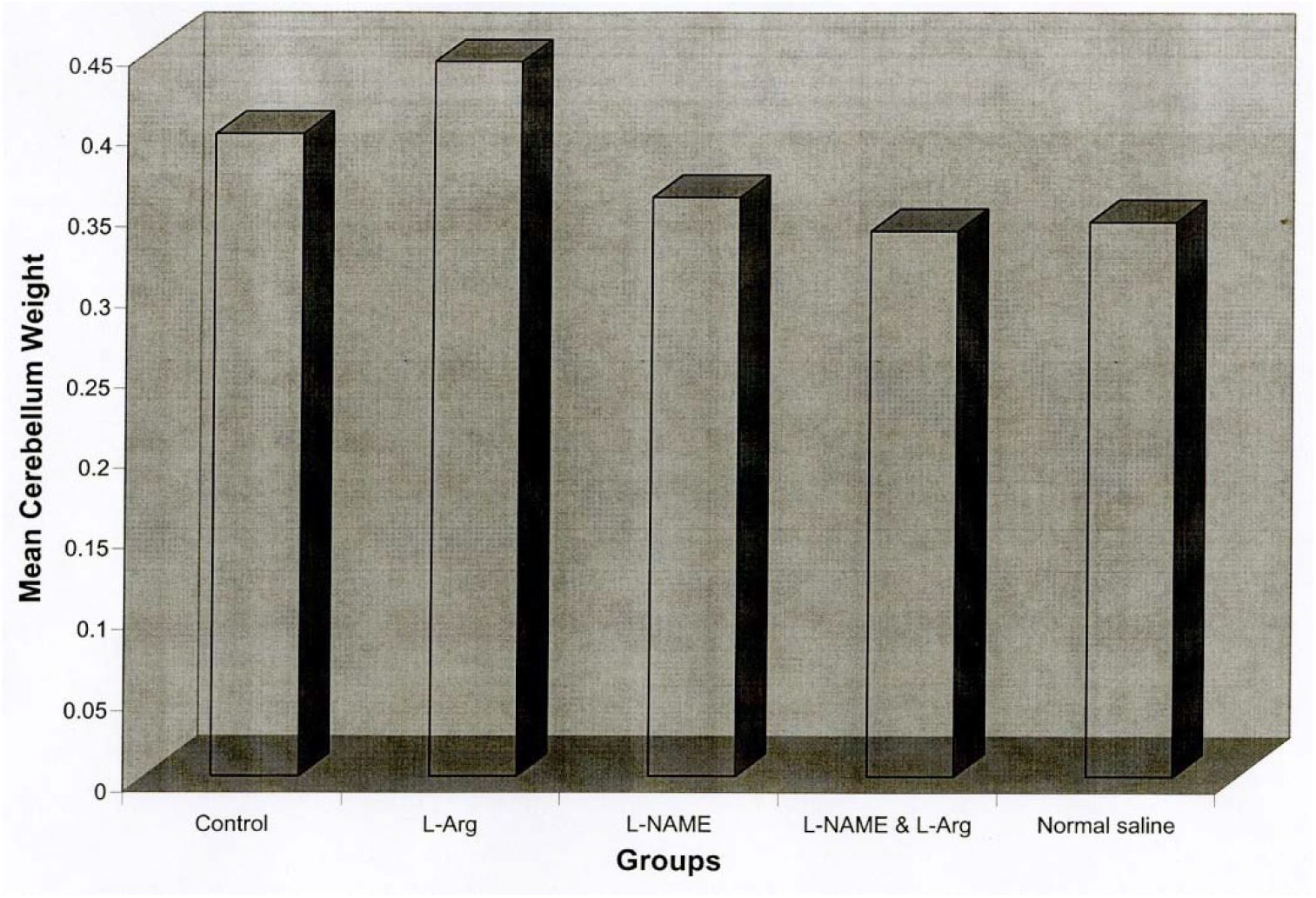
Bar chart of mean cerebellar weight in 5 control groups: L-Arg, L-NAME, L-Arg + L-NAME and normal saline.

**Table 4c.**
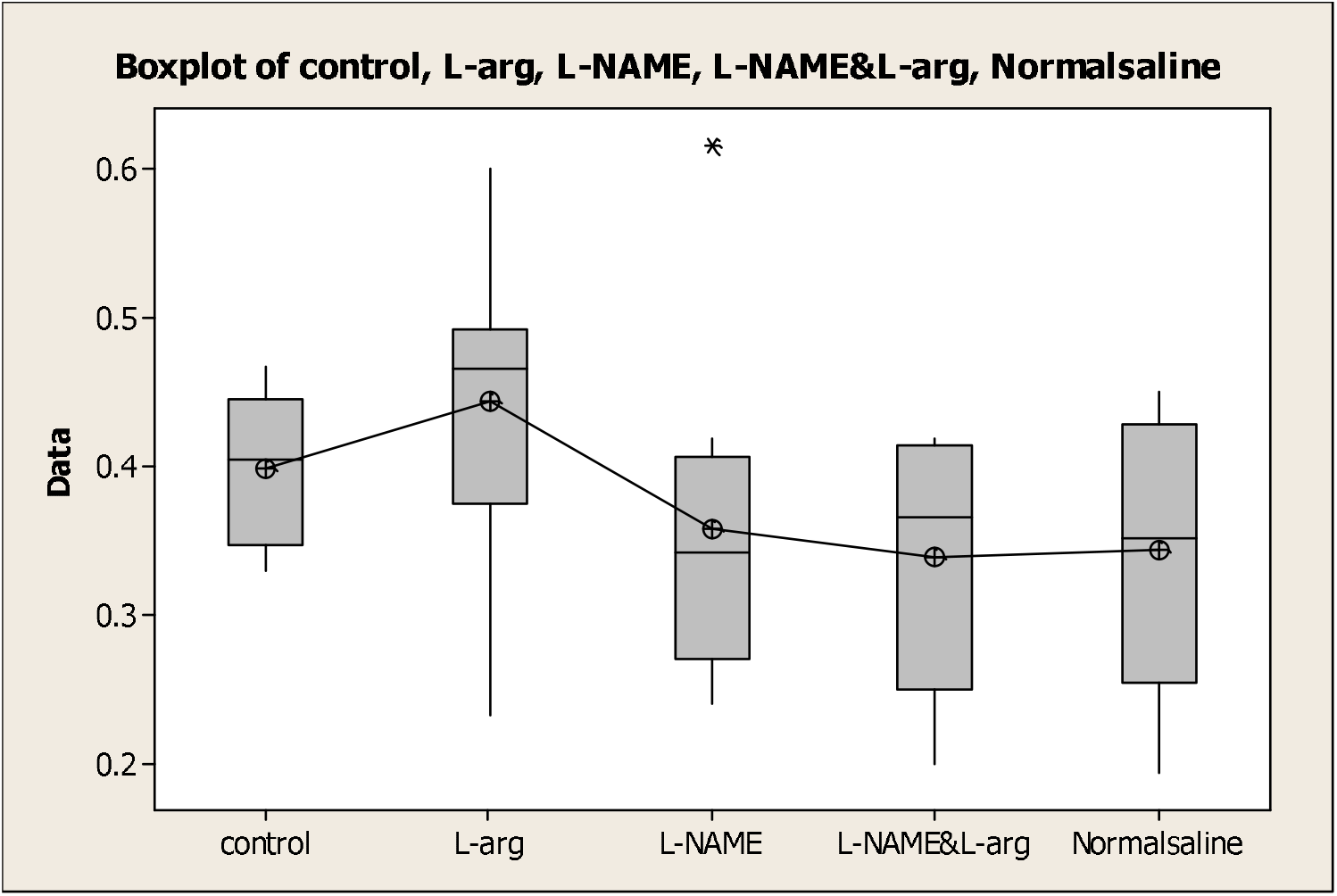
Boxplot graph of cerebellar weight in control, L-Arg, L-NAME, L-Arg + L-NAME and normal saline.

### Cerebellum volume

One-way analysis of variance was used to determine the relationship between the groups, which showed a statistically significant difference (P <0.01) between the groups. In the initial study of the groups with Tokey criterion at the level of 0.05 (P <0.01), a significant difference was observed between the L-arg group and the normal saline and L-NAME and L-NAME + L-Arg groups.

**Table 4a:** Quantitative values including mean and standard deviation of cerebellar volume in different groups. To the right is the 95% confidence interval of the mean..

| Individual 95% CIs For Mean Based on |  |  |  |
| --- | --- | --- | --- |
| Pooled StDev |  |  |  |
| Level | N | Mean | StDev |
| control | 8 | 0.5200 | 0.0447 |
| L-arg | 8 | 0.7143 | 0.1773 |
| L-NAME | 8 | 0.3125 | 0.1356 |
| L-NAME&L-arg | 8 | 0.3800 | 0.0837 |
| Normalsaline | 8 | 0.4200 | 0.1924 |

Pooled StDev = 0.1416

**Table 4b1:**
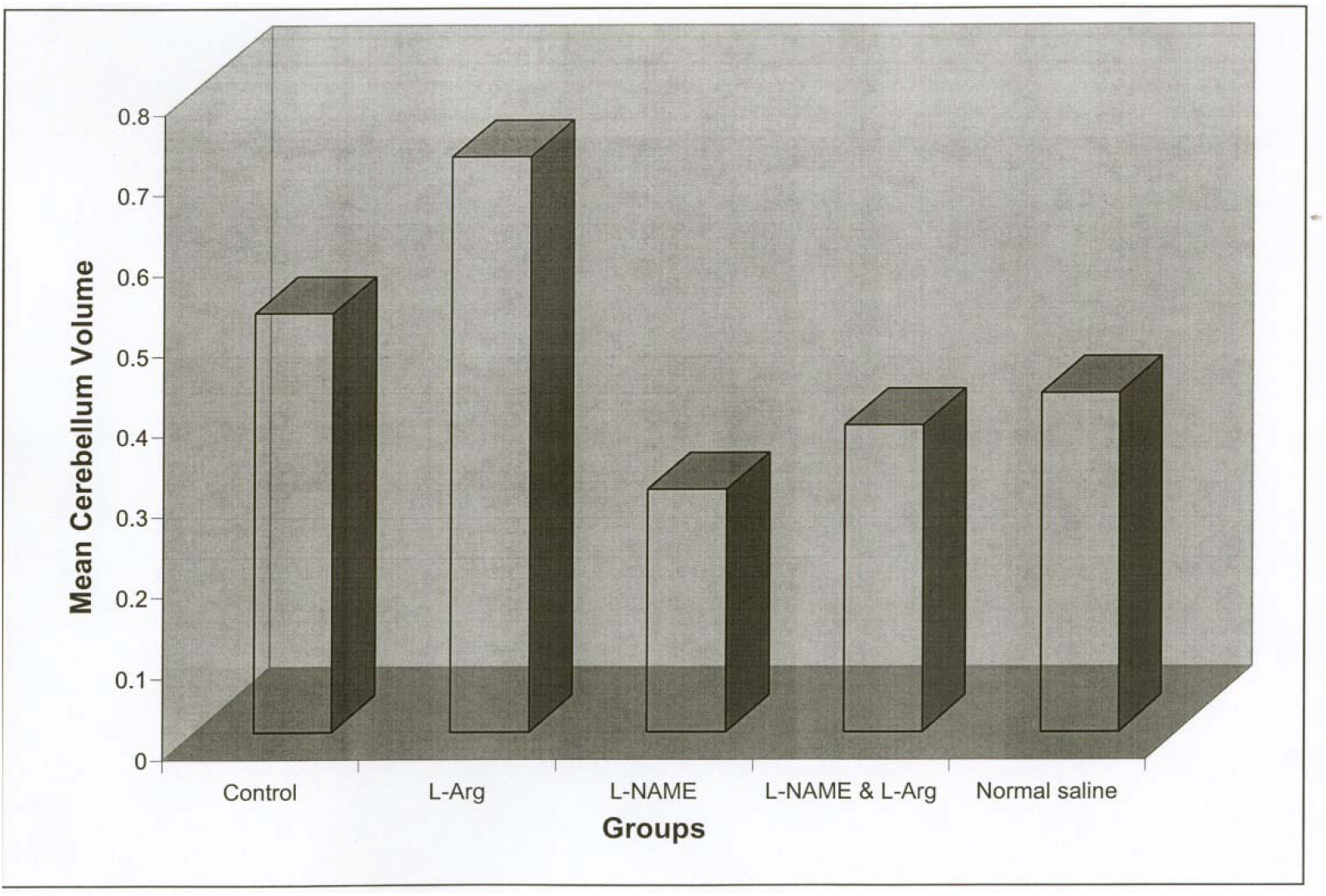
Bar graph of mean cerebellar volume in 5 control groups: L-Arg, L-NAME, L-Arg + L-NAME and normal saline.

**Table 4c.**
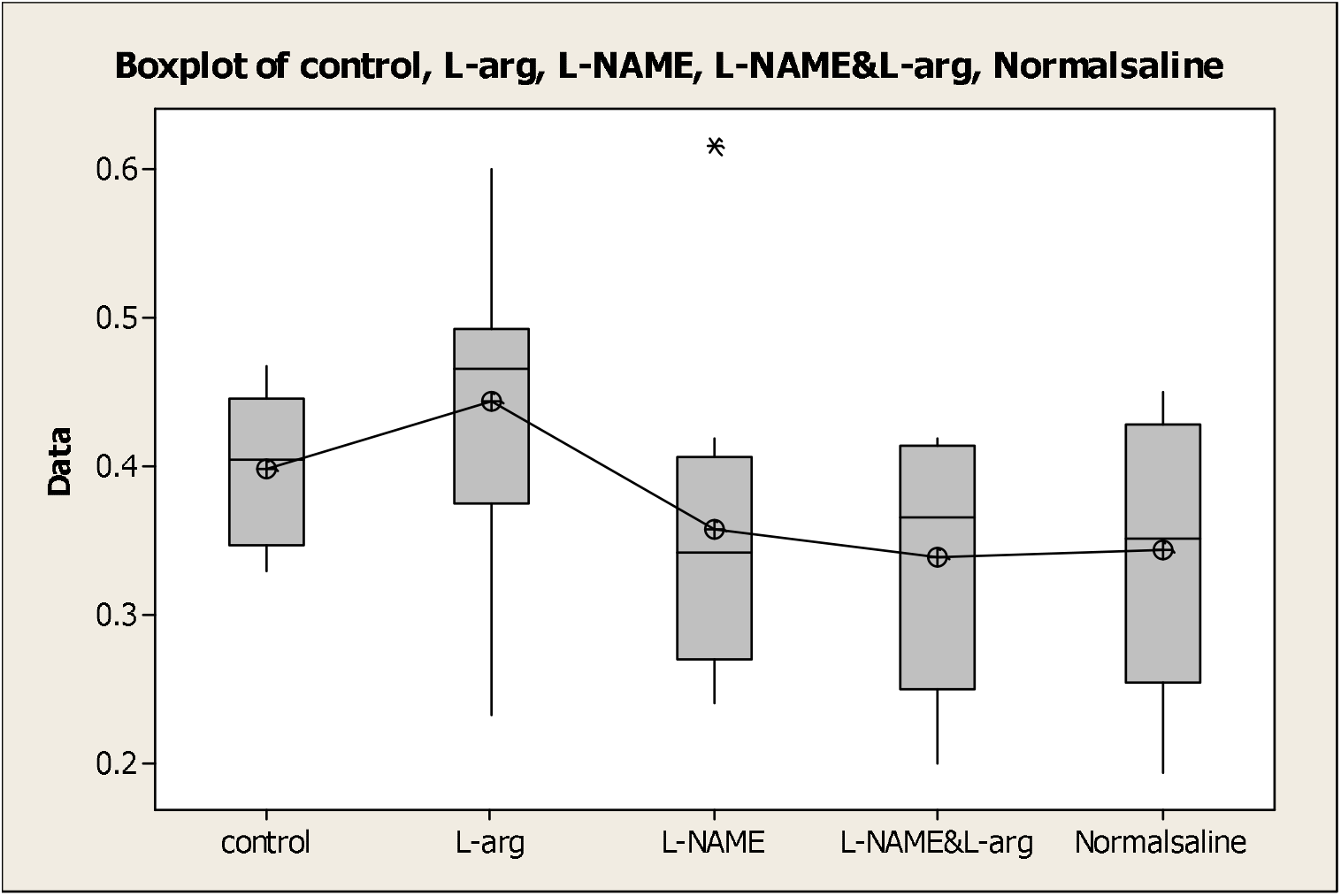
Boxplot diagram of cerebellum volume in control groups, L-Arg, L-NAME, L-Arg + L-NAME and normal saline.

But there was a significant difference between the control group and other groups and also between L-NAME, L-NAME + L-arg and normal saline groups.

### Microscopic results

In the microscopic study of the cerebellar slides of rats, the observations are as follows:

The cerebellar structure of rats in the normal saline group was not significantly different from the control group, The L-Arg + L-NAME group, on the other hand, underwent only minimal changes in cerebellar structure. In the L-Arg injected group, the granular and molecular layers were slightly thickened, some nuclei in the granular and molecular layer were severely hyperchromatized and Purkinje cells accumulation was seen with lymphocytic invasion. In the L-NAME group, the number of Purkinje cells seemed to have increased and a some of lymphocyte cells also invaded this accumulation (Photomicrographs 1 to 14).

**Figure 1 and 2:**
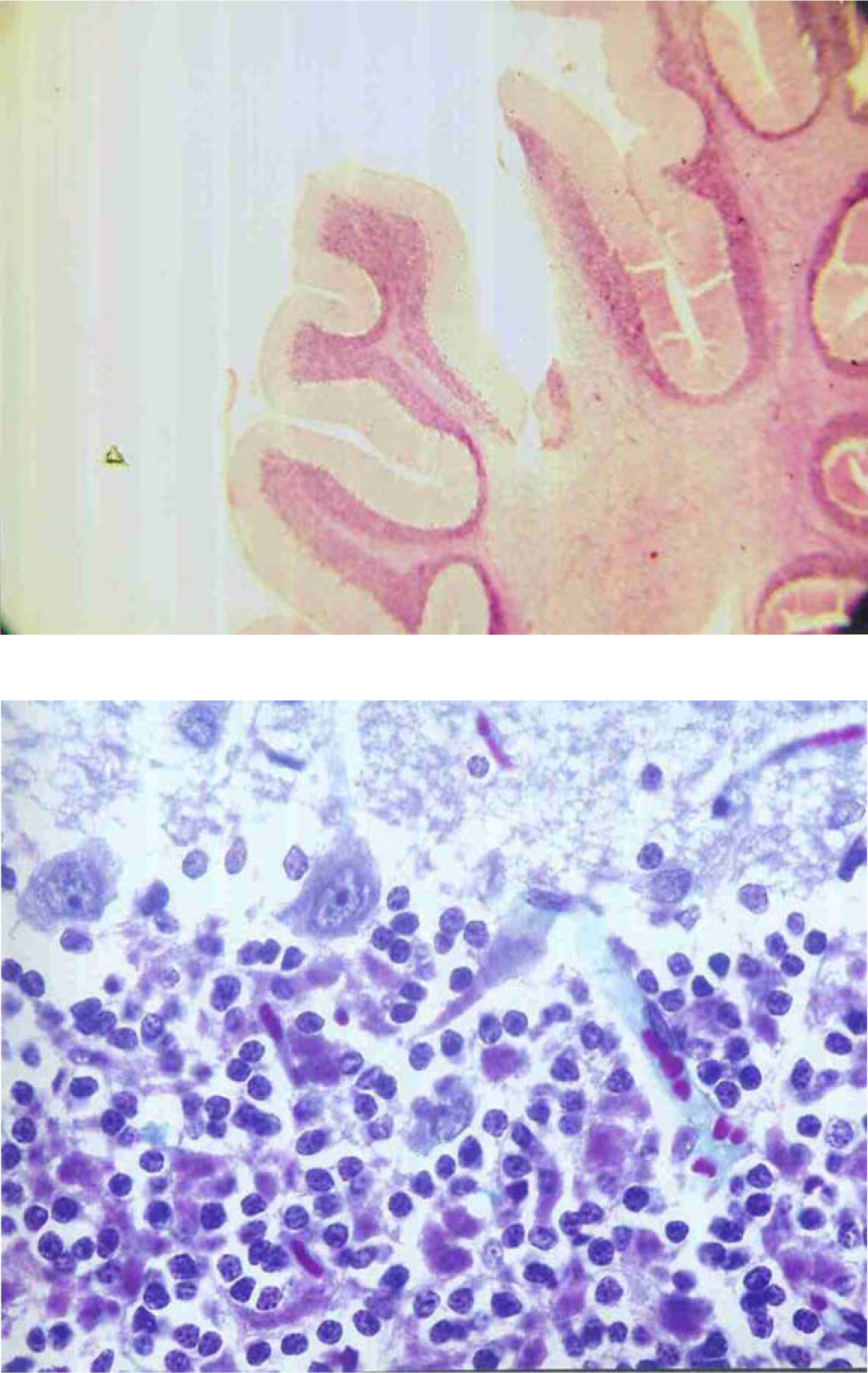
Cerebellar histological structure in the control group; photomicrograph 1, H & E staining with 4X and photomicrograph 2, Masson trichrome staining with 100X.

**Figure 3 and 4:**
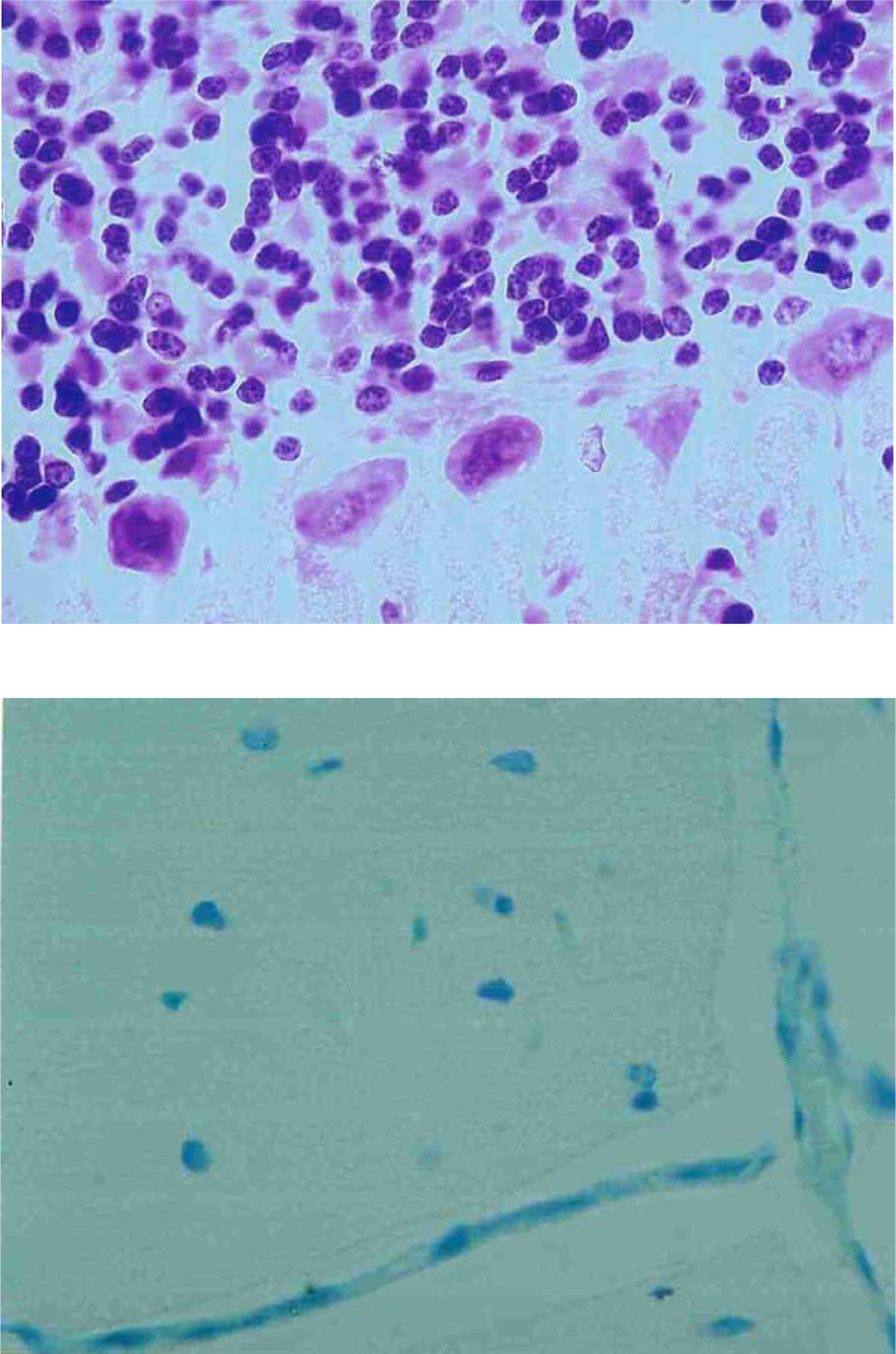
Cerebellar histological structure in the normal saline; As expected, no major changes are observed. Figure 3, H & E staining with 100X, indicates the Purkinje layer. Figure 4, Toluidine blue staining, indicates molecular layer with 100X.

**Figure 5 and 6:**
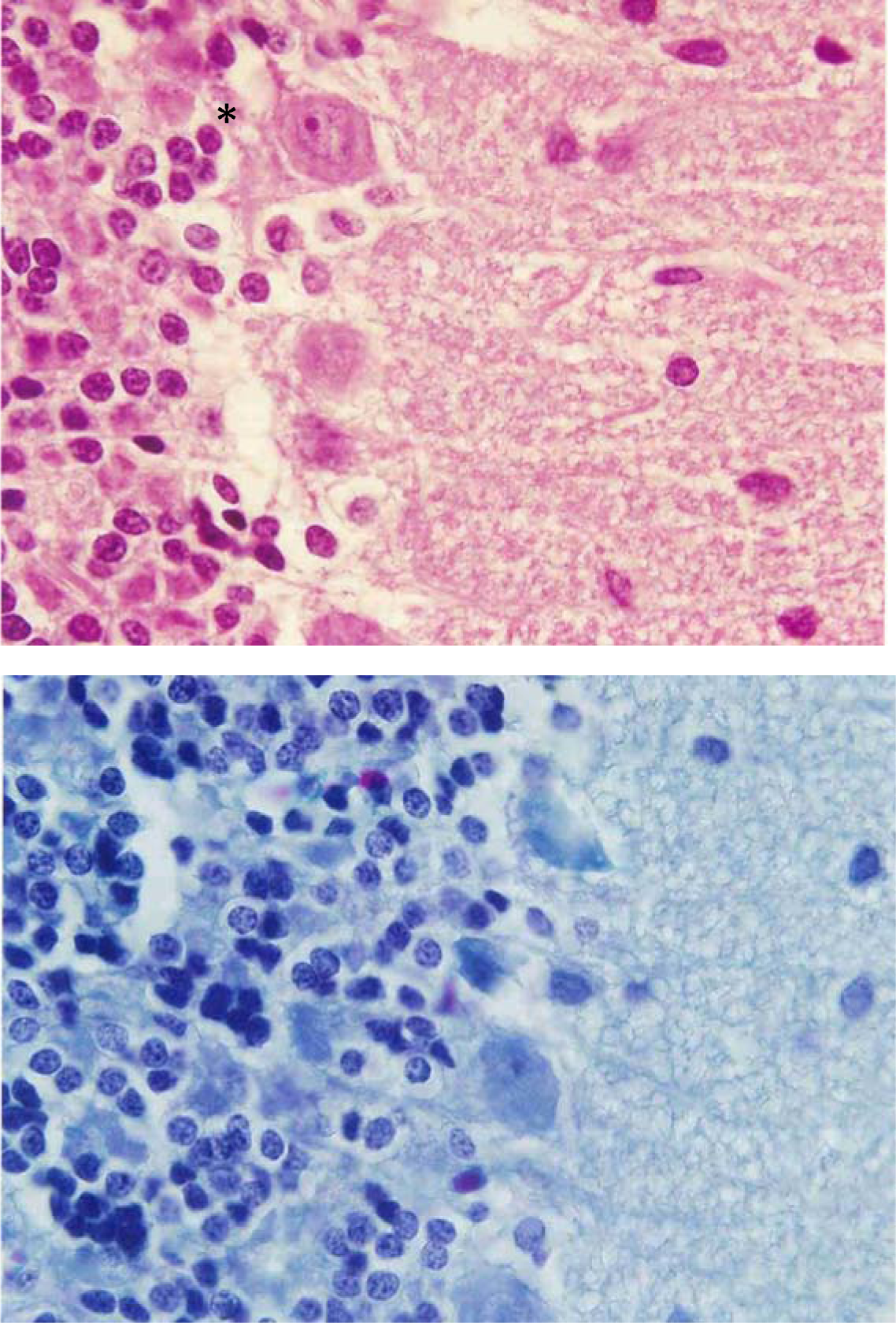
Cerebellar histological structure and changes in L-NAME group; Figure 5, taken with H&E staining with 100X, The number of Purkinje cells seems to have increased and some lymphocyte cells have invaded. Figure 6, taken with Mason trichrome staining with 100X, Numerous black dots indicate lymphocytic invasion.

**Figure 7,8 and 9:**
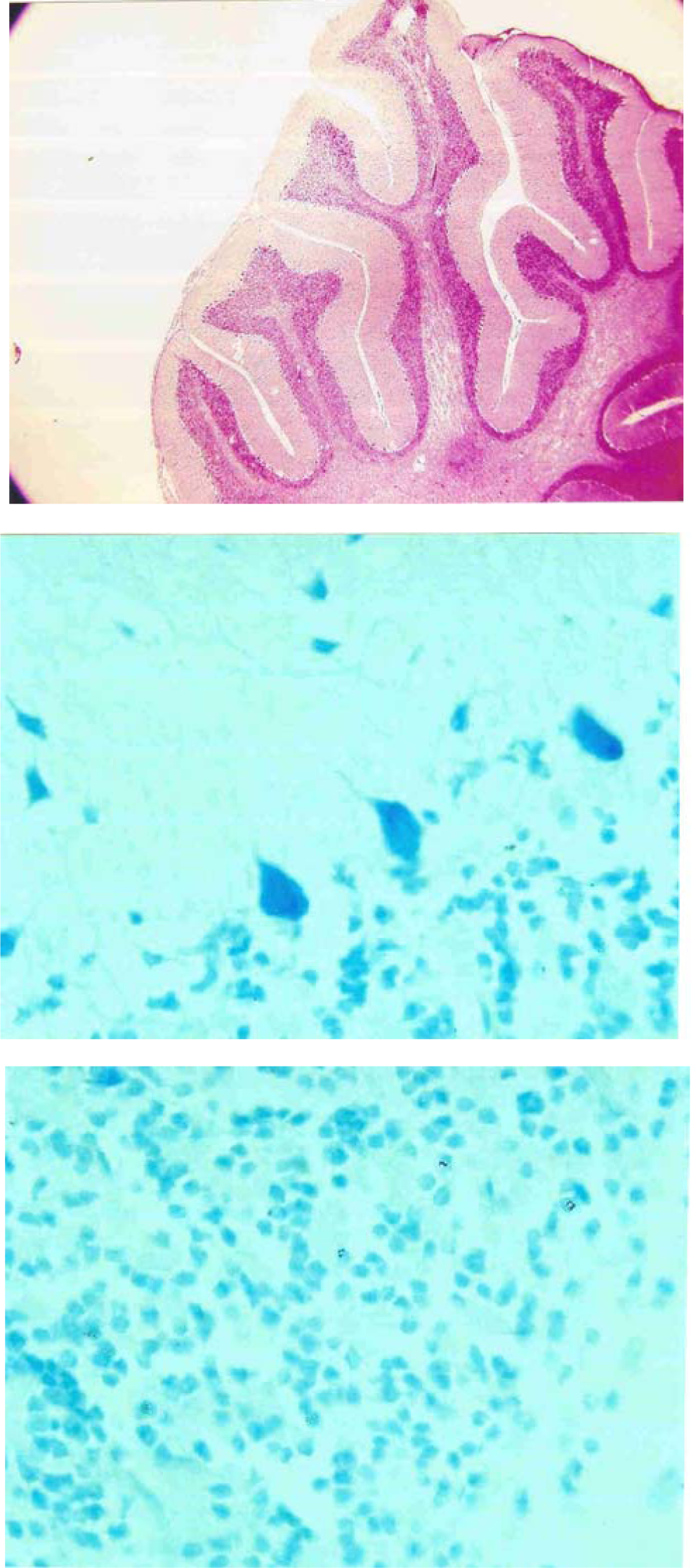
Cerebellar histological structure and changes in L-NAME + L-Arg group. Figure 7, taken with H&E staining with 4X; The opposite effects of these two substances have led to no significant change. Figure 8, Purkinje cell layer and Figure 9, Granular layer, both show with toluidine blue staining with 100X. The changes are negligible.

**Figure 10 and 11:**
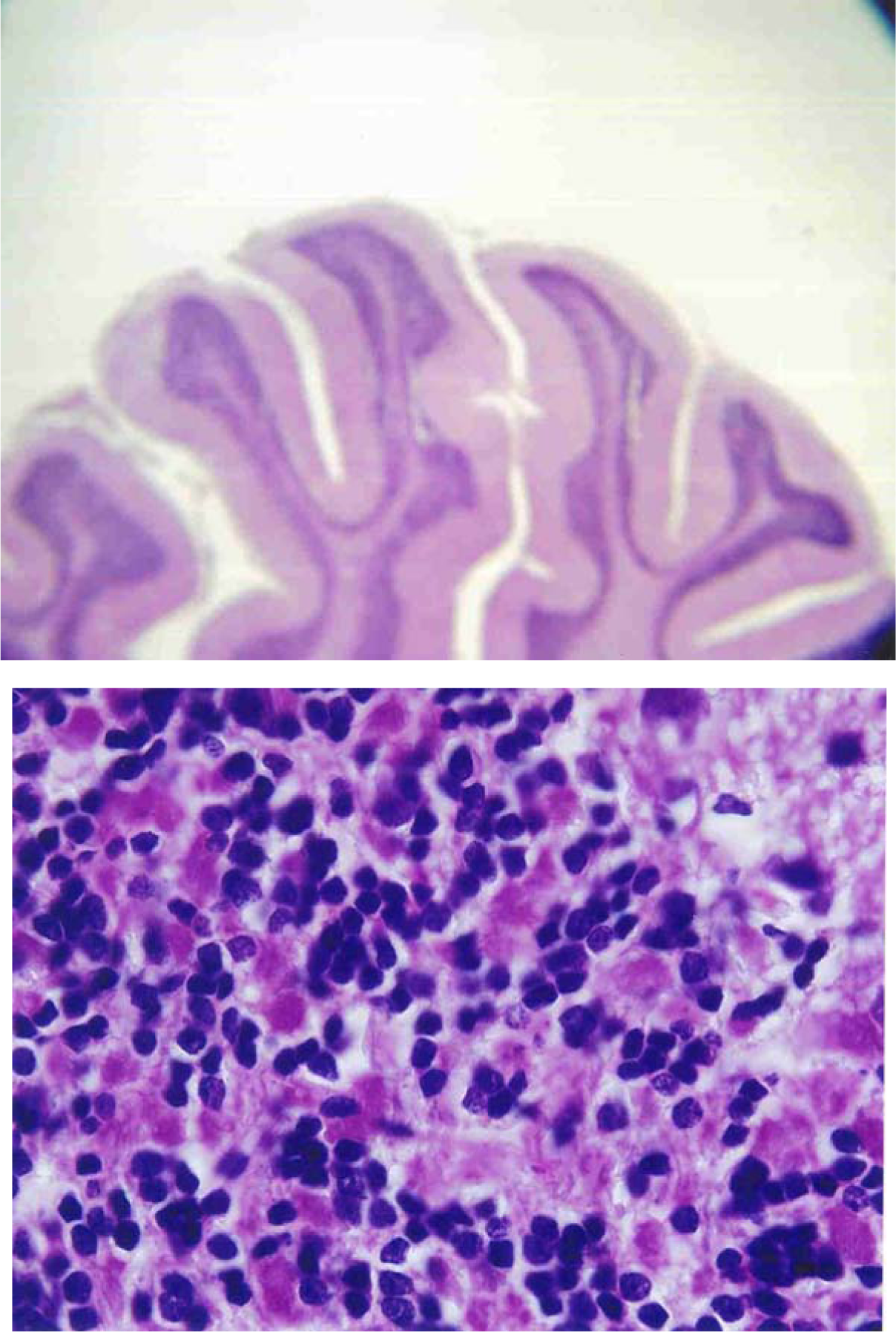
Cerebellar histological changes in L-Arg group. Figure 10 shows a slightly thicker granular and molecular layer (H&E staining with 4X); Figure 11 shows the presence of large amounts of acidophilic molecules in the granular layer Which is the extracellular matrix; Also the color differences of the nuclei show a severe hypochromatosis of some granular layer nuclei (H&E staining with 100X).

**Figure 12 and 13:**
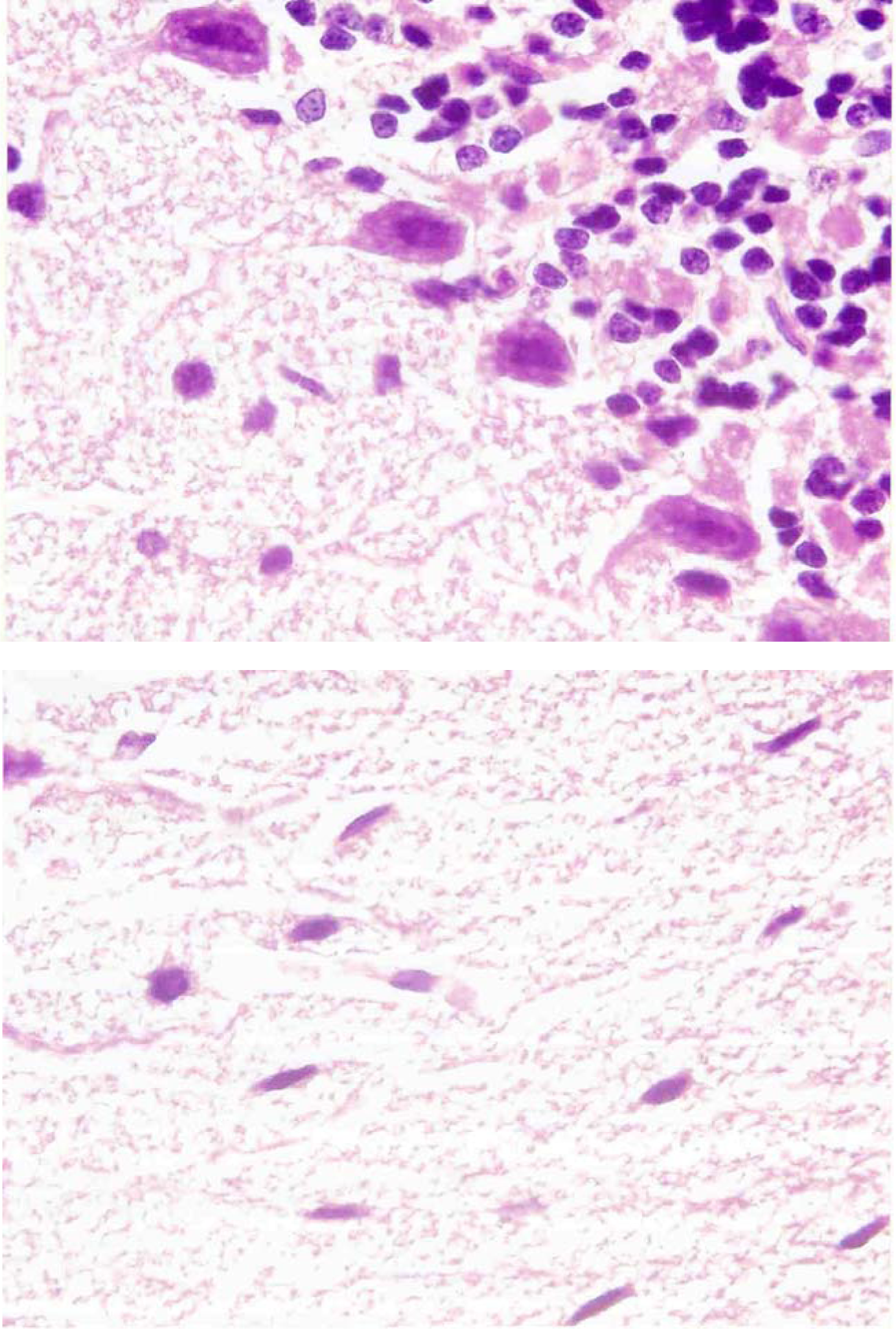
Cerebellar histological changes in L-Arg group. Figure 12, Purkinje cell proliferation and Figure 13, Molecular layer cell nucleus hyperchromatosis. The background of the molecular cortex is vesicular (Both figures, H&E staining with 100X).

**Figure 14:**
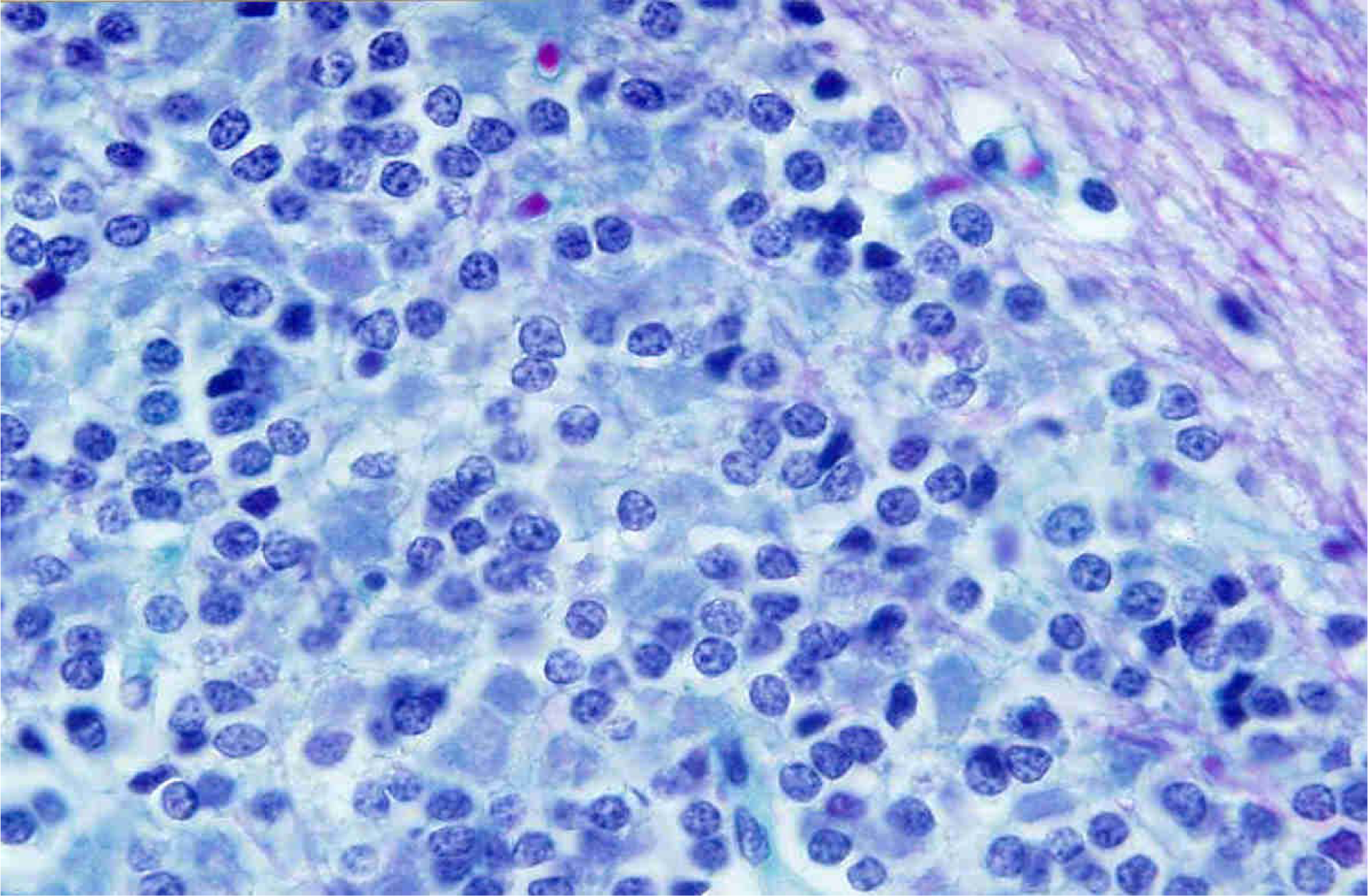
Cerebellar histological changes in L-Arg group. Accumulation of Purkinje cells is seen with lymphocytic invasion (Mason trichrome staining with 100X).

## Discussion

Nitric oxide (NO) is a simple diatomic molecule produced in the body of mammals and many physiological and pathological effects are attributed to this small molecule. The precursor to the in vivo production of this substance is L-Arginine and its synthesis is catalyzed by the NOS enzyme. NOS is inhibited by a substance called L-NAME. In this study, the quantitative and qualitative effects of nitric oxide on the histopathology of cerebellum by increasing or decreasing the in vivo production of this substance were investigated. In this study, forty Wistar female rats (RAT) with a weight of about 200 to 250 grams and an average age of eight weeks were used. Rats were divided into five groups of eight, including control groups, normal saline, L-NAME, L-Arginine, L-NAME + L-Arginine. On the third, fourth and fifth days, the injection was performed intraperitoneally and on the eighteenth day, after anesthesia with ether and then craniotomy, the brain and cerebellum of the animals were removed. After quantitative measurements, including weight and volume measurements, the organs were fixed in 10% formalin, and after performing tissue preparation steps for slide preparation, sections with a thickness of 5 to 6 microns were prepared and they were stained and evaluated by The general hematoxylin-eosin method and special technique such as Mason trichrome and toluidine blue.

### The effect of NO on the cerebellum

The results of this study suggest that NO precursors can increase microscopic blood flow to the cerebellum. In a group of rats injected with L-Arg, the granular and molecular layers were slightly thickened under the microscopic view; also Purkinje cell accumulations were observed, which showed a significant increase in volume as a macroscopic finding. This finding can be explained by the vasodilatory effect of NO and increased blood flow in the cerebellar microvascular. As Morikawa et al. demonstrated this vasodilating role by injecting L-Arg [11]. Also In Rodrigoez’s 2004 study, the role of NO as a vasodilator in the early stages of ischemia and cerebellar hypoxia was confirmed [13]. Lisa Mapelli et al. (2017) showed the granular layer in the cerebellar sections of rats which diameter of capillaries changes rapidly after stimulation of moss fibers, under the influence of nitric oxide Vasodilation requires stimulation of the NOS, which may occur in the pericytes. In this study, focusing on granular cells (GrC) of mossy fibers, it was shown that (GrC) activity enhances vasodilation of surrounding capillaries through the NMDA-nNOS receptor signaling pathway. As a result, these capillaries regulate localized blood flow according to the changes in neuronal activity in the cerebellum. These results show a signaling network that indicates the granular layer as an important determinant in regulating cerebellar blood flow [16].

In addition, in the microscopic view of this group, lymphocytic invasions were observed among Purkinje cells which indicate the inflammatory stimulatory effects of NO produced by NOS. This could explain the role of NO in many cerebellar pathologies. As Rodriguez also showed, if induction of NO synthesis continues following ischemic and cerebellar hypoxia, iNOS is activated and this time NO causes inflammatory reactions as a free radical [13]. Our study has shown this inflammatory response in the form of lymphocytic invasions.

Co-administration of L-NAME and L-Arg showed no inflammatory effects, which may be due to the inhibitory effect of L-NAME on NO production from L-Arg so it can be concluded that L-NAME and similar compounds can be used to reduce NO-induced tissue damage. This feature has been used in various studies to reduce the pathological complications of NO [9].

Another study in 2004 confirmed this neurotoxic role of NO from iNOS [14]. A study by Lin Wang et al found that the role of nitric oxide in neuronal development was challenging. In this study, cerebellar granular neurons (CGN) were used as a model to assess the role of nitric oxide in survival and related signaling pathways. Stable inhibition of nitric oxide production has been reported to cause apoptosis in differentiated granular neurons. The findings show that blockade of nitric oxide production induces apoptotic death through caspase-3 activation. This study provides direct evidence of the active role of nitric oxide in maintaining the survival of granular neurons [17]. L-NAME injection showed the proliferation of Purkinje cells (which does not lead to macroscopic changes) and lymphocyte accumulations, which can be explained by the effects of L-NAME on vascular stasis, capillary congestion [9], and also the effects of growth and inflammatory factors on congested vessels. Due to the extraordinary role of NO, it is suggested that the histopathological effects of this molecule in low and high doses on the brain and cerebellum in different complications, especially acute ischemia and hemorrhagic lesions, be studied and compared. Currently, only in cases of acute non-hemorrhagic cerebral ischemia, drug therapy (for instance TPA or tissue pelasminogen activator) is available; However, it seems that by intervening in the various effects of NO deficiency or abnormal rising, more effective preventive or therapeutic actions can be taken.

## List of abbreviations

Not applicable.

## Declarations

### Ethical considerations

all of the experiments which involve animals are ethically acceptable and conform to the guidelines for the care and use of laboratory animals.

### Consent for Publication

Not applicable.

### Availability of data and materials

Not applicable.

### Authors contributions

This article was designed and Supervised by Dr.Chegini and Dr.Noori. Ms.Chegini, Mr.Gholamzad, and Ms.Sadeghi participated in scientific writing and did Laboratory works that were analyzed by Dr.Chegini. All authors have read the manuscript.

## Acknowledgements

Not applicable.

